# Screening megasynthetase mutants at high throughput using droplet microfluidics

**DOI:** 10.1101/2023.01.13.523969

**Authors:** Farzaneh Pourmasoumi, Sundar Hengoju, Katharina Beck, Philipp Stephan, Lukas Klopfleisch, Maria Hoernke, Miriam A. Rosenbaum, Hajo Kries

## Abstract

Nonribosomal peptide synthetases (NRPSs) are giant enzymatic assembly lines that deliver many pharmaceutically valuable natural products, including antibiotics. As the search for new antibiotics motivates attempts to redesign nonribosomal metabolic pathways, more robust and rapid sorting and screening platforms are needed. Here, we establish a microfluidic platform that reliably detects production of the model nonribosomal peptide gramicidin S. The detection is based on calcein-filled sensor liposomes yielding increased fluorescence upon permeabilization. From a library of NRPS mutants, the sorting platform enriches the gramicidin S producer 14.5-fold, decreases the number of stop codons 250-fold, and generates enrichment factors correlating with enzyme activity. Screening for NRPS activity with a reliable non-binary sensor will enable more sophisticated structure-activity studies and new engineering applications in the future.

## Introduction

The shortage of effective antibiotics and the slow rate of antibiotics discovery have directed growing attention towards bioengineering of known antibiotic producers.^[1]^ A diverse portfolio of antibiotics and valuable natural products are synthesized nonribosomally in bacteria by non-ribosomal peptide synthetases (NRPSs).^[2,3]^ In NRPSs, chains of modules specifically incorporate one amino acid per module. In a minimal NRPS module, the adenylation (A-)domain selects and activates a specific substrate, the thiolation (T-)domain carries substrates between domains, and the condensation (C-)domain catalyses peptide bond formation. Modularity has made NRPSs a target for re-engineering^[4–6]^ although complexity and enormous sizes of the megasynthetases with ca. 130 kDa per module pose challenges. Previously reported tools for screening libraries of NRPS mutants only detect partial reactions or are limited in throughput.^[7–10]^ Therefore, a paucity of powerful high-throughput screening technologies remains an important bottleneck for re-engineering NRPSs.

Droplet-based microfluidics is a rapidly developing ultrahigh-throughput screening platform, allowing compartmentalization of molecules, cells, or library members into picolitre-volume reaction chambers, linking products to producers. ^[11–13]^ Diverse molecular properties have been screened for in microfluidics at ultrahigh throughput,^[14–19]^ but application on megasynthetases such as NRPSs remains elusive. Fluorescent reporter strains growing in microfluidic droplets together with potential producers of antibiotics have been employed to identify antibiotics in environmental samples^[20]^ or libraries of ribosomally produced and posttranslationally-modified peptides.^[19]^ However, these reporter strains remain notoriously low in sensitivity, suffer from the capriciousness of living organisms, and are prone to false positives caused by failure to grow or to generate the fluorophore. Sensor liposomes have been proposed as a robust alternative for detecting membrane-active antimicrobial peptides produced in droplets, where the antibiotic-induced permeabilization of fluorophore-filled liposomes generates a fluorescence signal.^[18,21]^

Here, we apply sensor liposomes encapsulating the self-quenching fluorescent dye calcein in microfluidic droplets to detect nonribosomal production of the membrane-active antimicrobial peptide gramicidin S (GS). Using this platform, we retrieved heterologously GS producing *Escherichia coli* cells from a 1:1000 dilution. Then, we performed site-directed mutagenesis on two positions in the GS NRPS to study structure-activity relationships and sorted the variants. Microfluidic droplet sorting of antibiotic producers from megasynthetase libraries will be a useful tool in the ongoing battle against antimicrobial resistance.

## Results

### Heterologous production of gramicidin S

To establish a new procedure for screening and selection of megasynthetase mutants, we chose the GS biosynthetic gene cluster as a model system. To allow genetic manipulation, the cluster has previously been cloned from the wild-type producer *Aneurinibacillus migulanus*^[7]^ and plasmid pSU18-grsTAB carrying the gene cluster has been constructed for production of GS in *E. coli* HM0079.^[22,23]^ The highest GS titer of 44 µM was achieved by incubation at 30°C and 300 rpm shaking for 48 h using buffered TB medium (Figure 1A). After centrifugation of the culture, triple quad LC-MS/MS analysis detected 90 % of the GS in the cell pellet and 10 % in the supernatant (Figure 1A). According to earlier reports, GS concentrations in this range can permeabilize bacterial^[24]^ or liposomal membranes.^[25]^

**Figure 1.**
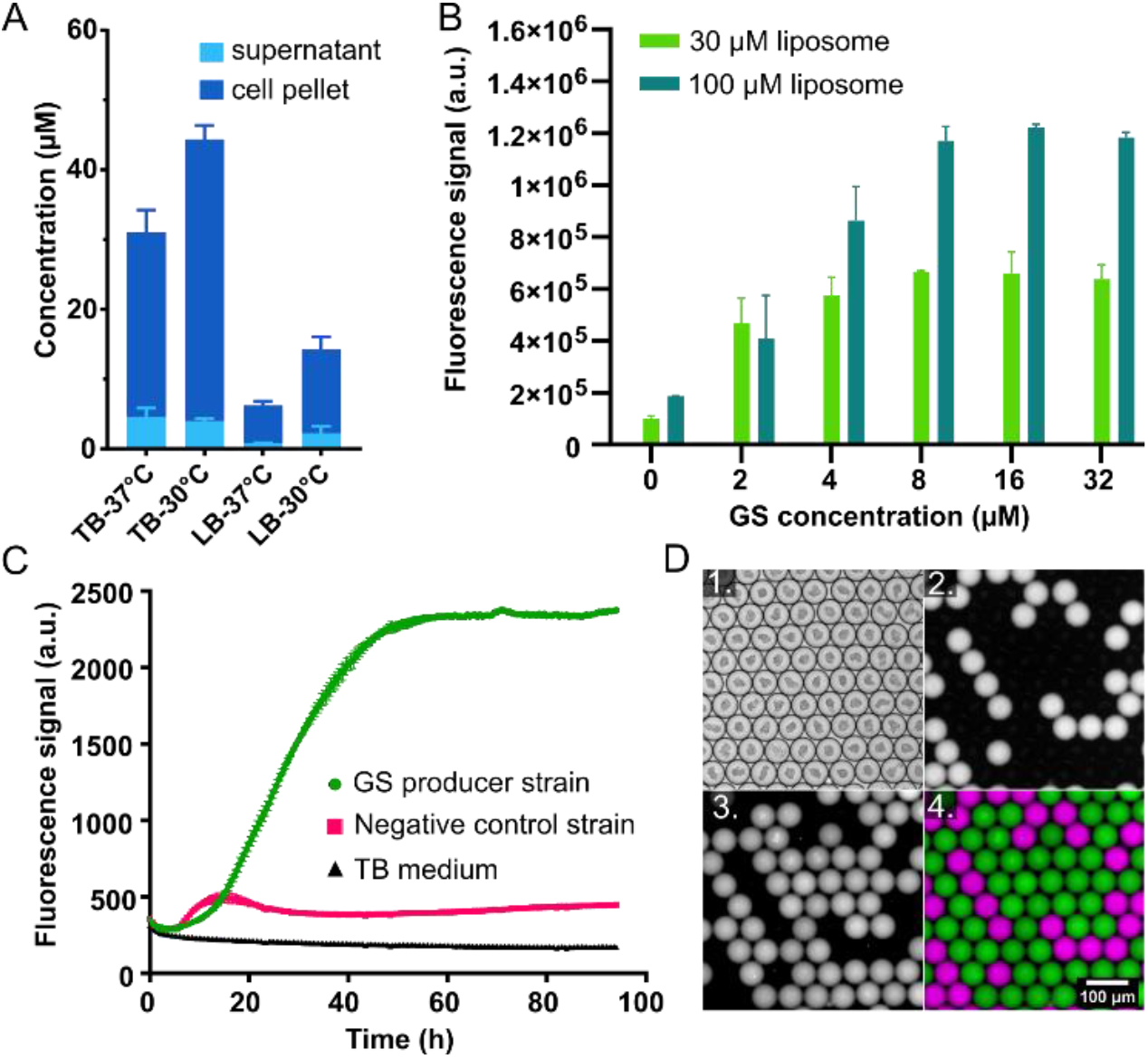
Detection of GS with a liposome sensor. **A**. Heterologous production of GS in *E. coli*. Error bars reflect three biological replicates. **B**. Detection of GS-induced permeabilization of liposomes in a microtiter plate. The concentration of POPC liposomes is given relative to the phospholipids. Error bars reflect two technical replicates. **C**. Detection of liposome permeabilization induced by GS produced by *E. coli*, the negative control strain without functional GS, and TB medium. Fluorescence indicating leakage is shown for three biological replicates. **D**. GS detection in droplets. Images of droplet populations containing liposomes initially filled with calcein and either *E. coli* that are GS producers as positive control or non-producers (labelled with red dye) as negative control. Images were taken at 10-fold magnification after 48 h incubation. 1. Bright-field, 2. Red fluorescence highlighting droplets with negative control strains, 3. Green fluorescence highlighting activation of the liposome sensor by GS, i.e. leakage of liposomes inside the droplets, 4. Overlay of red (colored in purple) and green fluorescence images.

### Liposome sensor for gramicidin S

To detect GS, we tested liposomes filled with calcein as a fluorescent sensor compatible with microfluidics.^[26]^ Calcein is self-quenching at the high concentrations entrapped inside the sensor liposomes. Through release and dilution of calcein into the droplet volume, fluorescence increases. For establishing the assay, GS was extracted and purified from the wild-type producer.^[22,27]^ First, we tested in microtiter plates whether purified GS would permeabilize calcein-filled liposomes within 30 min at concentrations between 2 and 32 µM GS (Figure 1B). With 30 µM liposomes, the fluorescence signal levels off already at 4 µM GS. With 100 µM liposomes (lipid concentration), an increase of the signal up to 8 µM GS was observed. The liposome sensor not only provides a strong signal over background but also a dynamic range rather than a binary signal. This dynamic range can be further adjusted through the liposome concentration inside the individual droplets. For our screening purposes, we used POPC liposomes (POPC = 1-palmitoyl-2-oleoyl-*sn*-phosphatidylcholine), but depending on the activity or lipid selectivity of the target molecule, tailored lipid compositions are also conceivable.^[26,28]^

### Detection of heterologous production of gramicidin S

Next, we validated the ability of the heterologous GS producer, *E. coli* HM0079::pSU18-grsTAB, to permeabilize liposomes in microtiter plates. As non-producing negative control, the same strain with a plasmid carrying only *grsA* was used.^[29]^ Only the GS producing *E. coli* strain triggered permeabilization of the sensor liposomes, but with medium or the negative control strain, liposomes were stable for at least 95 h (Figure 1C). Altogether, we found the sensor liposomes to be preferable over reporter strains for monitoring GS production because they are shelf-stable and reliable, provide a dynamic range of detection, and do not compete with the producer cells for resources.

### Detection of gramicidin S in microdroplets

For high-throughput screening and sorting, we implemented the liposome sensor in a droplet microfluidics setup (Figure 2A). This setup includes droplet generation, off-chip incubation, and fluorescence-activated sorting. *E. coli* cells and sensor-liposomes are encapsulated into water-in-oil droplets (~100 pL) by using a flow-focusing chip. The total GS titer produced inside droplets (23 µM) was lower than that obtained in a microtiter plate (44 µM), but still sufficient to effectively permeabilize liposomes. The fluorescence of calcein upon release from liposomes was stable and contained within the droplet confinement for at least 72 h (Figure S1). For bacterial growth and peptide production, droplets are incubated in a “droplet lung” providing homogeneous oxygen supply.^[30]^ Droplets with active expression of GS cause permeabilization of liposomes and release of fluorophore molecules resulting in high intensity of green fluorescence inside the droplet. For sorting, droplets are reinjected into a sorting chip, fluorescence intensities are measured using an optical fiber-based setup,^[31]^ and highly fluorescent droplets are collected for further analysis.

**Figure 2.**
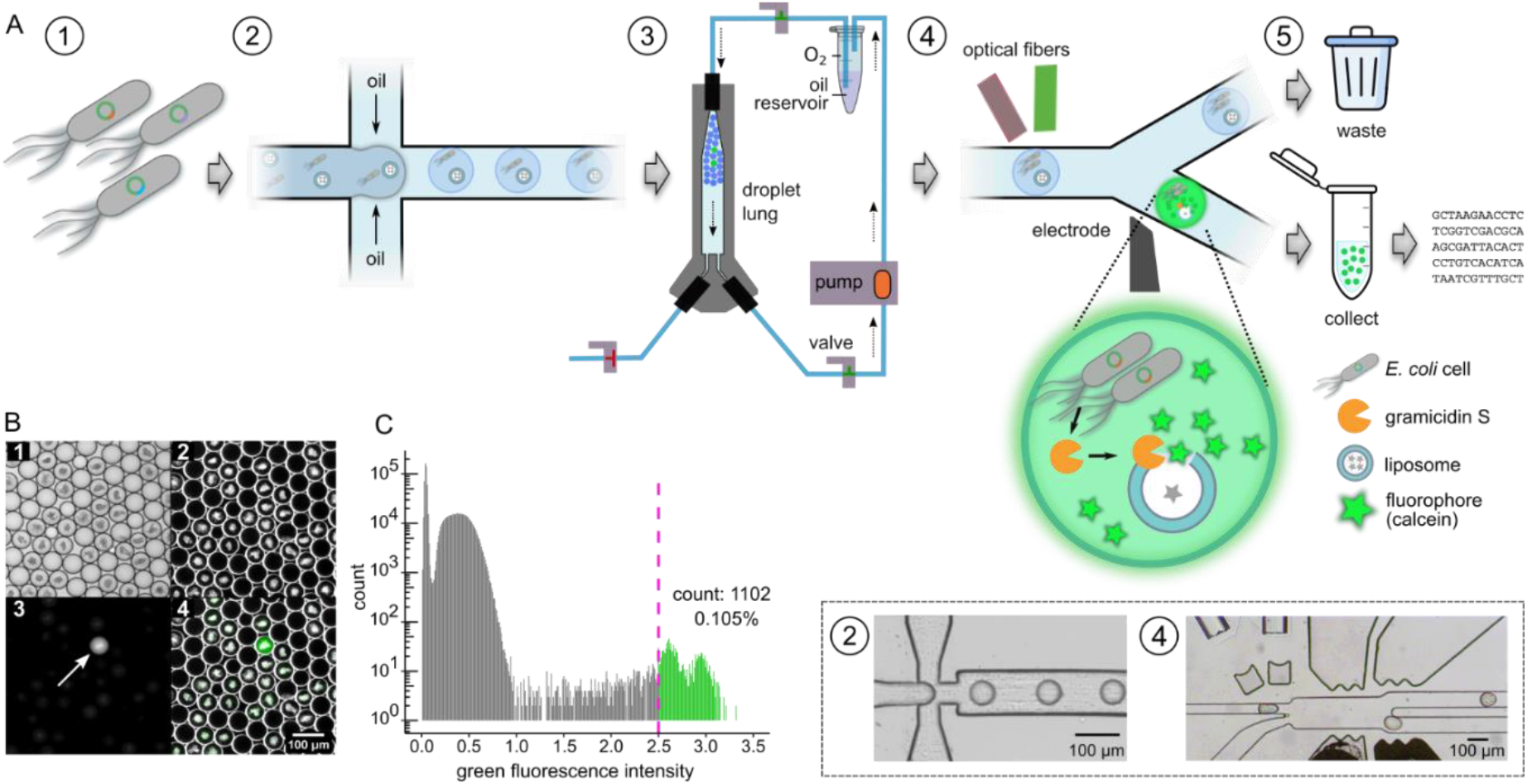
Fluorescence analysis and sorting of droplets. **A**. Schematic of the sorting platform. The experimental protocol consists of five steps. 1. Plasmid library preparation and transformation of *E. coli*. 2. Single-cell encapsulation into droplets. 3. Incubation in a droplet lung for *E. coli* growth and peptide production. 4. Fluorescence-triggered sorting of the library. 5. Pooling of the hits followed by plasmid purification and sequencing. Steps 2 and 4 are shown as microscopic images (box). **B**. A mixture of GS producing and non-producing *E. coli* cells are encapsulated at a 1 to 1000 ratio. For microscopic imaging, droplets are loaded into an observation chamber. Images show 1. bright-field, 2. dark-field, 3. green fluorescence channel, and 4. an overlay of dark field and green fluorescence channel. The arrow shows an example of a droplet containing the GS producer. **C**. Histogram of the green fluorescence intensity of droplets. The sorting threshold (pink dashed line) was set at 2.5 relative fluorescence units. Fluorescence droplets passing the sorting threshold were sorted, collected, and deposited on LB agar plates^[32]^ for further genetic analysis.

The microfluidic assay was validated on a mixture of two separately generated droplet populations both containing sensor liposomes (100 µM lipid concentration) and bacterial growth medium. The first droplet population contained GS producing *E. coli* cells, and the second population contained the negative control *E. coli* strain and a red fluorescence dye to tell them apart. Subsequently, a mixture of both droplet populations was incubated at 28°C for 48 h in the droplet lung. After incubation, fluorescence microscopy of droplets in an observation chamber showed strong green fluorescence in GS producer droplets, but not in negative control droplets (Figure 1D and S3).

### Sorting for gramicidin S production

To evaluate the sorting power of our microfluidics setup, we mixed GS producing *E. coli* cells directly with negative control *E. coli* cells at a ratio of 1:1000 and encapsulated the mixture into one droplet population. The cell density at the time of droplet generation was adjusted so that most droplets would contain zero or one cell according to a Poisson distribution with an average occupancy of ~36 %. After incubation in the droplet lung, a sample was analyzed with a microscope (Figure 2B). With the remaining droplets, fluorescence sorting was performed to retrieve those with the top 0.1 % of fluorescence intensity (Figure 2C). Utilizing an integrated droplet plating platform,^[32]^ 28 droplets from the sorted population of 1102 droplets were plated on LB agar supplemented with chloramphenicol. As control, another experiment was performed with deactivated sorting gate and 28 droplets from this average population were also plated. Genetic characterization of the extracted plasmids from both populations showed an increase from 0/28 GS producers without sorting to 28/28 after sorting with no detectable false positives. Compared to the initial ratio of 0.1 % GS producer cells, these counts indicate a 1000-fold enrichment factor. We conclude that the liposome sensor allows highly efficient retrieval of producer droplets even from a dilute population resembling the composition of genetic libraries typically used in directed evolution experiments.^[12]^

### Screening a library of megasynthetase mutants

With the screening platform established, we went on to sort a genetic library of the GS biosynthetic gene cluster. Mutations were targeted to an A-domain of the NRPS. A-domains act as “gatekeepers” and primarily dictate the sequence of peptide products.^[33–37]^ Two positions in the binding pocket of the GrsB1 A-domain (Figure 3A) were simultaneously mutated using degenerate NNK codons (Table S1). We chose positions I729 and A731 (Table S2) because these are located in direct vicinity of the substrate and thus part of the “specificity code”^[34,35]^ (Figure 3B, Table S3) but not too strictly conserved. An alignment with related Pro-A-domains (Table S3) showed some variation at these positions. Therefore, we expected to obtain a broad range of activity levels from fully active to fully inactive. To generate a large enough library despite the large size of plasmid pSU18-grsTAB (20 kb) harboring the GS gene cluster, the cloning strategy was carefully optimized. For more efficient transformation, we first created the library in a small helper plasmid only carrying the relevant *grsB* region. Next, the library fragment was transferred to pSU18-grsTAB, which was used to transform the production strain *E. coli* HM0079. Illumina MiSeq sequencing of the unsorted library confirmed that 85.5 % of the theoretically possible 400 side chain combinations were present in the library.

**Figure 3.**
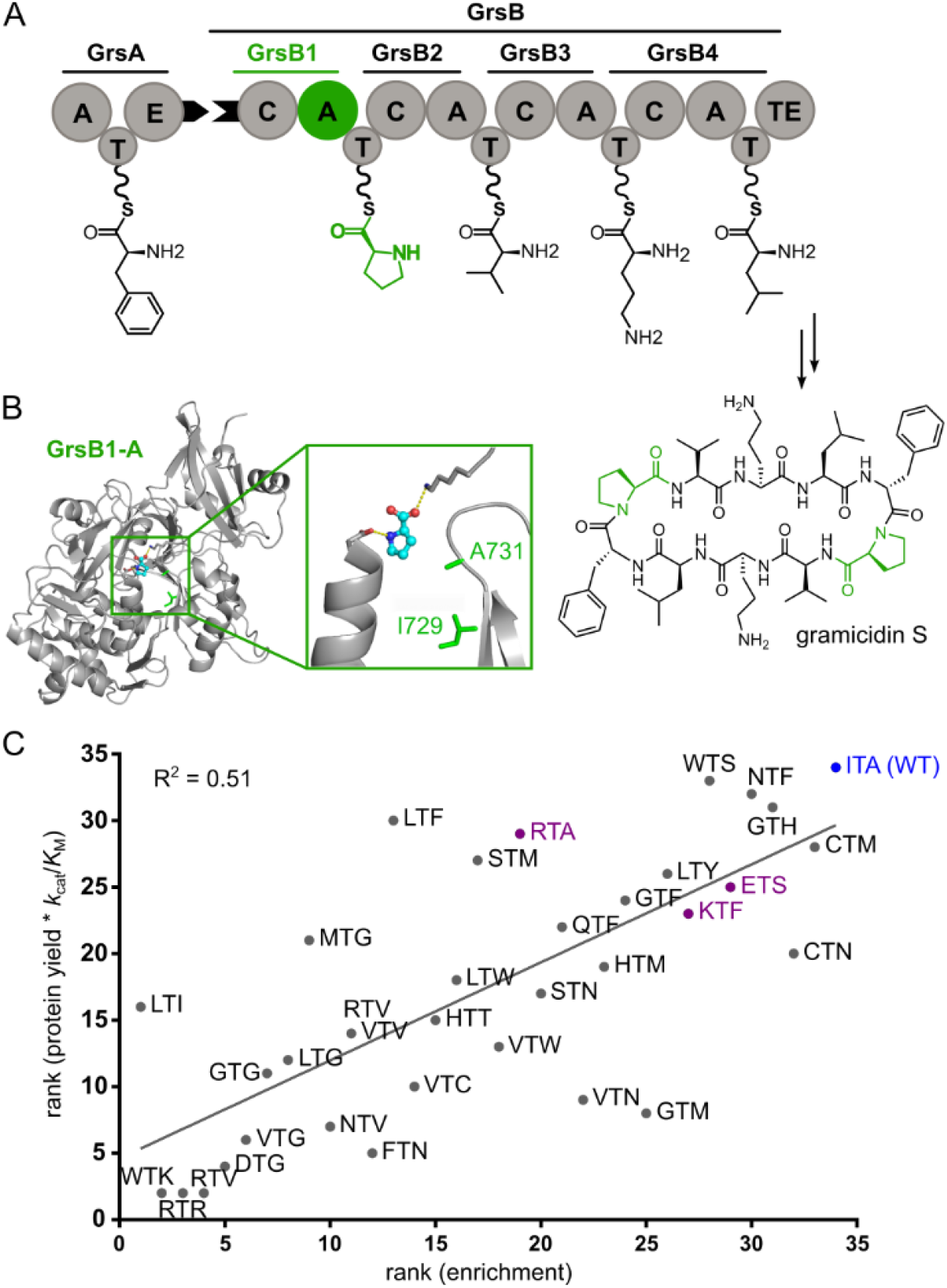
Validation of GS library screening results. **A**. Megaenzymes GrsA and GrsB synthesize GS.^[33]^ **B**. Homology model of the GrsB1 A-domain. The protein sequence of the GrsB1 A-domain was submitted to the SWISS-MODEL server for homology modelling^[40]^ and the Pro ligand was added by hand. The structural model of the A-domain was visualized using PyMOL. **C**. Spearman rank correlation of the *k*_cat_/*K*_M_ value for adenylation of Pro (measured by MESG/hydroxylamine assay) multiplied by the protein yield and the microfluidic enrichment factor (R^2^ = 0.51). The wild type is highlighted in blue. Three sequence variants (ETS, KTF, and RTA; in purple) were selected for further evaluation because charged residues were introduced.

Next, we screened the mutant library using the microfluidics platform. In total, 4.9·10^5^ droplets with an average occupancy of ~10 % were screened and sorted based on their fluorescence signal to select the brightest 0.2 % containing the most active producers (Figure S5). To sample the unsorted library, the same experiment was repeated with deactivated sorting gate. Plasmid DNA was recovered from the sorted and unsorted droplet populations. Then, the region of the plasmid carrying the mutations was amplified by PCR and the amplicon sequenced using the Illumina MiSeq method. Both the sorted and unsorted library yielded 3·10^5^ analyzable sequences. After sorting, stop codons in the library positions were depleted 250-fold and the wild-type sequence (ITA) was enriched 14.5-fold, indicating an excellent performance of the screening method. While in general, more conservative changes were favored over less conservative ones (underrepresentation of Arg, Trp, Pro), some mutations introducing charges were unexpectedly drastic, such as ETS, KTF, or RTA.

### Validation of screening results

To evaluate how the microfluidic sorting depends on the NRPS activity, we measured the enzyme activity of GrsB1 variants *in vitro*. Based on the enrichment from the unsorted to the sorted library, we selected 32 variants in addition to the wild-type covering the full range from strong depletion to strong enrichment (Table S4). We investigated the effect of each set of mutations on the adenylation activity by measuring saturation kinetics of hydroxylamine-stimulated pyrophosphate release with Pro as a substrate.^[38]^ The Spearman rank showed a correlation between adenylation efficiency (*k*cat/*K*M) and enrichment factor, which slightly improved when the *k*cat/*K*M was multiplied with the protein yield obtained for the GrsB1 variant (R² = 0.51, Figure 3C, Table S4). Importantly, this result indicates that the screening method can not only distinguish active and inactive NRPSs, but also levels in between. Using a hydroxamate assay for adenylation specificity with proteinogenic amino acids,^[39]^ we observed that all variants with detectable activity have preserved their proline specificity (Table S4). Also, for the strongly enriched mutants with charged residues in the active sites (ETS, KTF, RTA), the presence of adenylation activity was confirmed.

## Conclusion

In this work, we present an efficient setup for screening megasynthetase mutants at high throughput using a liposome sensor on a microfluidics platform. By directly detecting the membrane permeabilization caused by the antimicrobial peptide GS, we eliminate the problems associated with biological reporter strains, such as low sensitivity and high rates of false positives. Using calcein filled liposomes provides a strong, dose-dependent turn-on signal which makes the sorting process extremely reliable. Hence, false positives were undetectable after a single round of sorting with a positive control strain diluted 1:1000 in non-producers. Sorting of a site-directed saturation mutagenesis library of GS producers designed to cover a broad range of activities showed that the liposome sensor offers not only a binary distinction of active from inactive variants, but a gradual response. It has been demonstrated by others that a liposome sensor can even distinguish activity against different membrane compositions.^[18]^

While the liposome sensor is limited in scope to the detection of membrane-damaging peptides, the direct detection of bioactivity instead of screening for partial reactions involved in the biosynthesis is an advantage. It may or may not be possible to improve the pharmacological properties of gramicidin S or tyrocidine^[41]^ by modifying their sequence. However, other membrane-active molecules such as the polymyxins are life-saving drugs in the clinic despite their toxicity. Improving polymyxins would be a formidable goal^[42,43]^ and can now be put into practice using the membrane activity of the molecule. To fully realize the potential of our screening platform, reliable protocols for cloning large megasynthetase libraries will be crucial.

In summary, we present a microfluidics platform capable of retrieving active mutants from large libraries of megasynthetase mutants that are synthesizing membrane-active molecules. This platform has demonstrated excellent sorting precision and may find use in the design and engineering of nonribosomal pathways towards antibiotics.

## Acknowledgements

The authors thank Martin Roth and Lisa Mahler for technical assistance and advice, professor Donald Hilvert (ETH Zurich) for kindly providing the *E. coli* HM0079::pSU18-GrsA strain, and Maximilian Müll for assistance with HAMA measurements. We are thankful for the financial support of the Jena School for Microbial Communication (JSMC), Deutscher Akademischer Austauschdienst (DAAD), Deutsche Forschungsgemeinschaft (DFG, German Research Foundation) with Project ID 441781663 (HK and LK) and Project ID 415894560 (MH and KB), the Daimler und Benz Stiftung (MH and HK), and the Fonds der Chemischen Industrie.

