## Supporting Information for "Screening megasynthetase mutants at high throughput using droplet microfluidics"

---

[a] F. Pourmasoumi, P. Stephan, L. Klopfleisch, Dr. H. Kries
Junior Research Group Biosynthetic Design of Natural Products
Leibniz Institute for Natural Product Research and Infection Biology (HKI)
Beutenbergstraße 11a, 07745 Jena, Germany
*

[b] Dr. S. Hengoju, Prof. Dr. M. A. Rosenbaum
Bio Pilot Plant
Leibniz Institute for Natural Product Research and Infection Biology (HKI)
Beutenbergstraße 11a, 07745 Jena, Germany
*

[c] K. Beck, Dr. M. Hoernke
Faculty of Chemistry and Pharmacy
Albert Ludwigs University of Freiburg, 79104 Freiburg i. Br., Germany

[d] Prof. Dr. M. A. Rosenbaum
Faculty of Biological Sciences
Friedrich Schiller University Jena, 07743 Jena, Germany

### These authors contributed equally.

### Supplementary Methods

#### Cloning

Inserts and vectors were assembled using In-Fusion cloning (Takara Bio) for transformation of *E. coli* Stellar cells. In-Fusion cloning was performed following the supplier's instructions. Primers for PCR amplification of inserts (Table S1) were designed with 15-20 bp long overlaps complementary to the insertion position in the vector DNA and ordered as synthetic oligonucleotides (Eurofins GmbH). Competent cells were transformed by heat shock. All inserts were checked by Sanger sequencing (Genewiz). Restriction enzymes for linearization of vectors were purchased from New England Biolabs (NEB). PCR reactions were performed with Phusion or Q5 High-Fidelity polymerase (NEB).

**Construction of pTrc99a-grsAB1_EcoNI.** The large size of plasmid pSU18-grsTAB makes handling during cloning difficult, therefore we created the smaller plasmid pTrc99a-grsAB1_EcoNI as a helper plasmid by PCR amplification of the relevant *grsB* region between two EcoNI sites on pSU18-grsTAB[1] followed by In-Fusion cloning of the PCR product in linearized plasmid pTrc99a. The *grsB* fragment was amplified from pSU18(F)-grsTAB with primers pTrc99a_GrsAB1_EcoNI_fw, and pTrc99a_GrsAB1_EcoNI_rev (Table S1). The vector pTrc99a (commercially available, GenBank: M22744, U13872, A13038) was linearized with XhoI and BamHI restriction enzymes (Figure S1).

**Construction of pTrc99a-grsTAB_nnk helper plasmid.** The GrsB1-A domain NNK Library was cloned by assembling two overlapping fragments harboring NNK codons at positions I729 and A731. Fragments A and B were amplified from pTrc99a-GrsB1 with primer sets GrsB1_AflII_fw / GrsB1_I729_rev and GrsB1_I729nnk_A731nnk_fw / pTrc99a_GrsAB1_EcoNI_rev and were cloned into pTrc99a-grsAB1_EcoNI linearized with AflII and BamHI via In-Fusion cloning (Figure S4). The plasmid library was purified from an overnight culture.

**Construction of pSU18-grsTAB_nnk Library.** Plasmid pSU18(F)-grsTAB was linearized with EcoNI. The fragment containing the NNK codons was amplified from the pTrc99a-grsTAB_nnk helper plasmid using primers GrsAB1_grsTAB_fw and GrsAB1_grsTAB_rev and was assembled with EcoNI linearized pSU18-grsTAB following the Infusion cloning protocol. The resulting plasmid library was purified from *E. coli* Stellar cells and re-transformed into *E. coli* HM0079 competent cells using heat shock transformation. The *E. coli* strain HM0079 (GT869 nrdD::sfp Spcr) carries a genomically integrated *sfp* gene encoding an unspecific phosphopantetheine transferase, which is essential for the production of NRPSs.[2]

**Site directed mutagenesis.** Selected variants from the sorted library were re-cloned for protein production. To make cloning of libraries into plasmid pTrc99a-GrsB1 more efficient, an artificial KpnI restriction side was introduced at codon V969 through silent mutation. Plasmid pTrc99a-grsB1 was first digested with AflII and BamHI. Two overlapping PCR amplicons created with the primer pairs pSU18-GrsB1_V969KpnI_fw / pSU18-GrsB1_BamHI_rev and pSU18-GrsB1_V969_rev / GrsB1_AflII_fw, respectively, were cloned into the linearized vector by means of In-Fusion cloning. The GrsB1-A library was cloned by assembling two overlapping fragments harboring suitable codons at positions I729 and A731. Fragments A and B were amplified from pTrc99a-GrsB1_EcoNI with primers sets GrsB1_AflII_fw / GrsB1_I728_rev and GrsB1_KpnI_fw_XXX / GrsB1_KpnI_rev, where XXX represents specific codons for library members on individual forward primers (Table S1), and were cloned into pTrc99a-grsB1_KpnI linearized with AflII and BamHI via In-Fusion cloning. For protein expression, the resulting plasmids were purified from *E. coli* Stellar cells and re-transformed into expression strain *E. coli* HM0079 competent cell using heat shock transformation.

#### Protein expression and purification

The *E. coli* strain HM0079 (GT869 nrdD::sfp Spcr)[2] was used for protein production if not stated otherwise. For heterologous expression of proteins in *E. coli*, a preculture was prepared by inoculating a single colony of the desired strain in 3 mL LB medium supplemented with the appropriate antibiotic followed by overnight incubation at 37 °C with 230 rpm shaking. Of the preculture, 500 µL was used to inoculate 400 mL of liquid 2YT medium containing appropriate antibiotics in a 2 L Erlenmeyer flask. The main culture was incubated at 37 °C, 230 rpm until the OD600 reached 1.0. At this point, the temperature was reduced to 18 °C and after 30 min further incubation, a final concentration of 250 µM isopropyl-β-D-thiogalactopyranosid (IPTG, Thermo Fisher Scientific) was added followed by incubation overnight at 18 °C, 230 rpm. Cells were harvested by centrifugation at 6,000 g for 10 min at 6 °C. The cell pellet was resuspended in 25 mL of low imidazole buffer (50 mM Tris-HCl, 0.5 M NaCl, 20 mM imidazole, 2 mM TCEP, pH 7.4) followed by addition of 100 µL of protease inhibition cocktail (P8849, Sigma-Aldrich). The cell suspension was sonicated on ice for 5 min at an amplitude of 40% in 2 s intervals with 3 s breaks with a UP200St ultrasonic processor (Hielscher). The cell lysate was centrifuged at 19,000 g for 30 min at 4 °C. The supernatant from the cell lysate was added to an Econo column (Bio-Rad) loaded with 2 mL of Ni-IDA beads (Rotigarose-His/Ni Beads, Carl Roth) and equilibrated with low imidazole buffer. The desired $His_6$-tagged protein was eluted in 4 fractions of 750 µL of high imidazole buffer (50 mM Tris-HCl, 0.5 M NaCl, 300 mM imidazole, 2 mM TCEP, pH 7.4) and samples from the last three elution steps were pooled and collected. Eluted protein was then buffer exchanged into protein storage buffer (50 mM Tris-HCl, 200 mM NaCl, pH 7.5) by centrifugation at 4,000 g and 4 °C in Vivaspin filters (Sartorius) with a molecular weight cut-off threefold smaller than the size of the protein. The protein concentration was calculated from the OD280 measured using a Take3 Plate on an Epoch2 microplate spectrophotometer (BioTek Instruments) with calculated extinction coefficients. For storage, 5% glycerol (Carl Roth) was added to the protein samples, which were flash frozen in liquid nitrogen and stored at -80 °C until use. Protein purity was analyzed by SDS-PAGE using Bolt 4-12% Bis-Tris Plus Gels (Fisher Scientific) with MES-SDS running buffer. Sample was loaded in Bolt LDS sample buffer supplied with Bolt reducing agent. Triple Color Protein Standard lll (Serva) was run alongside the protein samples as a size standard. The gels were run at 200 V for 22 minutes and stained with Quick Coomassie stain (Serva).

#### Kinetic evaluation via MESG assay

Michaelis-Menten parameters of the adenylation reaction catalysed by all selected library members were determined from kinetic data recorded with the MesG/hydroxylamine assay. The assay was performed as described previously with minor modifications.[3] Reactions contained 50 mM TRIS (pH 7.6), 5 mM $MgCl_2$, 100 µM 7-methylthioguanosine (MesG), 150 mM hydroxylamine (adjusted to pH 7.5-8 with NaOH), 5 mM ATP (A2383, Sigma), 1 mM TCEP, 0.4 U/mL inorganic pyrophosphatase (I1643, Sigma), 1 U/mL of purine nucleoside phosphorylase from microorganisms (N8264, Sigma) and varying amounts of TycA (0.025 – 1 mM) and substrates. In flat-bottom 384-well plates (781620, Brand) 100 µL reactions were started by addition of substrate and the absorbance was followed at 355 nm on a Synergy H1 (BioTek) microplate reader at 30°C. Background activity was recorded in wells containing buffer without substrate and the obtained slopes were subsequently subtracted. Each substrate concentration was measured in duplicate. Initial velocities were divided by the slope of a pyrophosphate calibration curve to obtain the pyrophosphate release rate. Initial velocities v0/[E0] were fit to the Michaelis-Menten equation by nonlinear regression using R version 3.4.2.

#### Quantification of gramicidin S by UPLC-MS/MS

Quantification of GS was performed on a Waters ACQUITY H-class UPLC system coupled to a Xevo TQ-S micro (Waters) tandem quadrupole instrument. The injection volume was 2 µL. The flow rate was 0.5 mL $min^{-1}$. Acetonitrile (B) and water with 0.1% formic acid (A) were used as strong and weak eluent, respectively. Acetonitrile was used to wash the injection needle between samples. Data acquisition and quantification were done using the MassLynx software (version 4.1). MS/MS analyses were performed using an ESI source in positive ion mode. Nitrogen was used as desolvation gas and argon as collision gas. The following source parameters were used: capillary voltage 0.5 kV, desolvation temperature 600 °C, desolvation gas flow 1000 L $h^{-1}$.

Column: ACQUITY UPLC CSH C18, 1.7 µm particle size, 2.1 × 50 mm
Elution profile: linear gradient of 40 to 98% B over 1 min followed by 1.2 min re-equilibration.
MRM transition: 571.6957 > 70.0991
Calibration curve range: 0.05 µM to 10 µM
Cone voltage: 16 V

#### POPC liposome preparation

Liposomes were prepared as described in the supporting information of Shi *et al.*.[4] Briefly, 100 mg POPC lipid (1-palmitoyl-2-oleoyl-*sn*-glycero-3-phosphocholine, Avanti Polar Lipids, Alabaster, AL, USA) were dissolved in 3.3 mL chloroform. The stock solution was aliquoted into amber glass vials (1.1 mL in 4 mL vials). Chloroform was removed under a gentle stream of nitrogen, followed by lyophilization overnight. Dry lipid films were used immediately or stored at -20 °C. In the next step, the lipid film was hydrated with 1 mL calcein buffer (70 mM calcein, 0.5 mM EDTA, 25 mM MOPS, pH 7.5). The hydrated lipid suspension was vortexed until the lipid film was fully suspended. The resulting multilamellar vesicles (MLVs) were subjected to five freeze-thaw cycles. For one cycle, the solution was frozen in dry ice for 3 min and thawed in warm water (55°C) for 3 min. The suspension was then vortexed for 20 s. Large unilamellar vesicles (LUVs) were prepared by extrusion by 41 passes through a hand extruder (LiposoFast hand extruder; Avestin, Ottawa, Canada) equipped with a 100 nm filter following the manufacturers instruction (Nucleopore Track-Etched Membranes, Whatman International Ltd, Maidstone, UK). In the last step, the calcein buffer outside the vesicles was exchanged for iso-osmotic external buffer without fluorescent dye (130 mM NaCl, 0.5 mM EDTA, 25 mM MOPS, pH 7.5) using a PD-10 desalting column (volume 2 mL, GE Healthcare, Little Chalfont, UK). Fractions of two drops each were collected from the column. The relevant fractions were only slightly colored and were pooled. The final lipid concentration of the LUV solution was determined using the Bartlett assay[5] as described in the Supporting Information of ref. [4].

#### Hydroxamate assay

**Reaction conditions.** The hydroxamate assay (HAMA) for adenylation activity was performed as previously described.[6] Reactions were conducted at room temperature in 100 µl volume containing 50 mM TRIS (pH 7.6), 5 mM $MgCl_2$, 100 mM hydroxylamine (pH 7.5-8, adjusted with NaOH), 5 mM ATP (A2383, Sigma), 1 mM TCEP and 1 mM of proteinogenic amino acids in 100 mM TRIS (pH 8). Reactions were started by adding enzyme solutions to a final concentration of 0.5 µM for the WT and 1 µM for all mutated enzymes. Reactions were quenched after 1 h by adding 10 µl reaction mixture to 90 µl 95% acetonitrile in water containing 0.1 % formic acid and submitted to UPLC-MS/MS analysis.

**UPLC-MS/MS conditions for HAMA**. Chromatography was performed on a Waters ACQUITY H-class UPLC system (Waters) with an injection volume of 3 µL. Water with 0.1 % formic acid (A) and acetonitrile with 0.1 % formic acid (B) were used as strong and weak eluent, respectively. Amino acid hydroxamates were separated on the ACQUITY UPLC BEH Amide column (1.7 µm, 2.1 x 50 mm) with a linear gradient of 10-50% A over 5 min (flow rate 0.4 mL/min) followed by 4 min re-equilibration. Water containing 0.1% formic acid was used as a needle wash between the samples. Data acquisition and quantitation were done using the MassLynx and TargetLynx software (version 4.1). MS/MS analyses were performed on Xevo TQ-S micro (Waters) tandem quadrupole instrument with ESI ionisation source in positive ion mode. Nitrogen was used as a desolvation gas and argon as collision gas. The following source parameters were used: capillary voltage 1.5 kV, cone voltage 65 V, desolvation temperature 500°C, desolvation gas flow 1000 L/h. GS was detected via specific mass transitions recorded in multiple reaction monitoring (MRM) mode. Standard calibration solutions of hydroxamates were prepared ranging from 0.0032 to 10 µM. The only detectable hydroxamate was ProHA, for which the lower limit of detection was 0.08 µM.

#### Microtiter plate assay with bacterial strains

Pre-cultures of *E. coli* cells (GS producer and negative control) were cultured overnight in TB medium (3 mL) supplemented with 6 mM ornithine and 25 µg/ml chloramphenicol at 37 °C, 230 rpm agitation. The main cultures were prepared in 96 well plates with 200 µl culture medium per well and 3 replicates per condition. Each well was inoculated with the pre-culture (1:500 v/v). A set of media-containing wells were not inoculated with pre-cultures as a second type of negative control. Liposomes (100 µM) were added to all wells at the time of inoculation. The fluorescence (Ex: 467 nm, Em: 515 nm) and absorbance (600 nm) were measured for 95 h, at 28°C and 750 rpm agitation (Figure 1C).

#### Microfluidics

**Chip fabrication**. Microfluidic chips for droplet generation and sorting were fabricated from polydimethylsiloxane (PDMS) by using soft lithography from a glass mold based on previously described procedure.[7] In brief, the 3D designs of channel layout were created in AutoCAD 2015 (Autodesk Corp.) and then printed using femtosecond laser machining (FEMTOprint) to prepare glass molds. Replicas of intermediate and working chips were prepared by using respectively 5:1 and 10:1 mixture of PDMS base and curing agent (Sylgard 184, Dow Corning). PDMS mixture was poured onto the mold, degassed in a vacuum desiccator, and thermally polymerized at 70 °C for 3 h. The polymerized PDMS was peeled off from the mold and cut into individual chips. The inlet and outlet for fluidic and electrical connections were punched using micropunches (Ted Pella). Finally, PDMS chips were plasma-bonded (Zepto, Diener) on microscope glass slides (Carl Roth) for droplet generation, and ITO coated glass (Delta Technologies) for droplet sorting. All microfluidic channels were treated with 1% (vol/vol) trichloro(1H,1H,2H,2H-perfluorooctyl)silane (Sigma) in Novec HFE-7500 (3M). Optical fibers were inserted into sorting chips in a designated fiber guide channel for fluorescence detection.[8] Electrodes were made by inserting low melting indium alloy Indalloy 19 (Indium Corp.).

**Culture preparation.** Culture medium TB was prepared as described previously.[9] *E. coli* cells (GS producer and negative control) were cultured overnight in TB medium with 50 µg/ml chloramphenicol at 37 °C. From the overnight culture, a main culture was prepared with a starting optical density of 0.1 at 600 nm (OD) and grown to mid-exponential phase. The cell culture used for droplet generation was prepared by diluting the pre-grown main culture to an OD of 0.1 using fresh TB medium and adding liposomes (200 µM) immediately before the experiment. For mutant library samples, cells were grown overnight after transformation and diluted to an OD of 0.001 (for droplets of volume ~100 pL), which should result in zero or one cell per droplet upon encapsulation according to a Poisson distribution.

**Fluid actuation and operation.** Fluids were actuated using pressure pumps (MFCSTM-EZ, Fluigent) and high precision syringe pumps (neMESYS, Cetoni). Polytetrafluoroethylene tubings (0.5 mm internal diameter, SCP GmbH) were inserted into the inlets and outlets of the microfluidic chips for transferring fluids. Novec HFE-7500 (3M) with 2% (v/v) PicoSurf (Dolomite) was used as continuous phase during all microfluidic operations. Microfluidic operations were monitored with an inverted microscope (Axio Observer Z1, Zeiss) and a high-speed camera (acA1920-150uc, Basler). A flow-focusing structure was used to generate droplets with a volume of ~100 pL. Generated droplets were collected in a dynamic incubation chamber and incubated at 28 °C for 48 hrs. Droplets were imaged in a stationary observation chamber with a PCO.edge camera (PCO) and a light source (Spectra-X, Lumencor).

**Optical and electrical setup for sorting.** For sorting of droplets, a fluorescence based optical setup with triggering system (Figure S2) was operated with little modification from our previously explained setup.[10] In brief, the excitation light from the laser source (488 nm, LASOS) was guided by an optical fiber (LASOS, core diameter 6 µm, cladding diameter 125 µm, and numerical aperture [NA] = 0.14) into the microfluidic channel. The emitted fluorescence light from droplets was collected by a second optical fiber (Thorlabs, core diameter 105 µm, cladding diameter 125 µm, and NA = 0.22), filtered through a quad-bandpass filter (440/521/607/700 HC Quadband, AHF) and detected by a photomultiplier tube (H10721-20, Hamamatsu) connected to a lock-in amplifier (HF2LI, Zurich Instruments). The output signals were acquired, analysed, and visualized using a FPGA card (National Instruments) in conjugation with a custom-written LabVIEW program. Sorting gates were adjusted within the application to send trigger signals and sort desired droplets by applying an alternating electrical field through a function generator (33210A, Keysight) and a high voltage amplifier (Trek). The function generator was operated in a burst mode with 450 mV (1000x amplified), 100 cycles, 20 KHz, and 50 % duty cycle. Sorted droplets with high fluorescence intensity were either collected in an Eppendorf tube for plasmid isolation or stored in an on-chip collection chamber and later distributed on agar plates.[11]

**Image analysis.** Stationary droplet images were analysed for determining grey values of each droplet in different channels using the image analysis software Fiji.[12] Initially, the contrast of bright-field images was enhanced for binarization using the default threshold method with a radius of 10 pixels. The binary structures were then dilated in a lateral direction by 5 pixels and subsequently inverted. Circular droplet borders were detected as regions of interest (ROIs) by applying the "Analyze Particles" function with an area of 2,000 – 20,000 pixels and circularity above 0.9. After reducing the radii of ROIs by 5 pixels to remove interfering dark pixels from actual droplet borders, ROIs were used to determine average grey values for respective channels. Measured values were analysed and visualized in the statistical software R.[13]

**Plasmid recovery and preparation for sequencing.** Sorted droplets were pooled and the emulsion was broken using an antistatic gun. Plasmid DNA was isolated using the DNA Clean and Concentrator – 5 kit (Zymo Research). The desired region on GrsB1 was amplified by PCR (30 s at 98°C, 35 x [10 s at 98 °C, 20 s at 61°C, 20 s at 72°C], 120 s at 72°C, final hold at 4 °C) using GrsB1-N-seq-4 and GrsB1_H765_rev primer pair and Q5 High-Fidelity DNA Polymerase (New England Biolabs). The purified DNA fragments were sent for next generation sequencing (using the Illumina MiSeq method).

**Production level of gramicidin S in droplets.** To investigate the GS production level, we generated droplets containing the producer strain in TB medium in a microfluidic setup. After generation, droplets were collected and transferred into an "droplets lung" with homogeneous oxygen supply[9] and incubated for 48 h at 28°C, after which GS was extracted from a population of ~300K droplets by breaking the droplets. This sample was centrifuged (11,000 x g, 4 min, RT) and the supernatant removed. The cell pellet, which contains most of the GS, was resuspended in 70% ethanol, followed by sonication for 10 min and further incubation at 60°C for 30 min. The cell debris was removed by centrifugation (19,000 x g, 4 min, RT). The supernatant was further diluted 10-fold in 50% ethanol containing 0.1% formic acid and the concentration of GS was quantified using UPLC-MS/MS in comparison with a standard purified from *A. migulanus*.[1] Concentrations were calculated assuming the GS was homogenously distributed in all droplets. We measured a total GS titer of 23 µM, of which 6 µM was in the supernatant and 17 µM was detected in the cell pellet extract.

#### Supplementary Figures

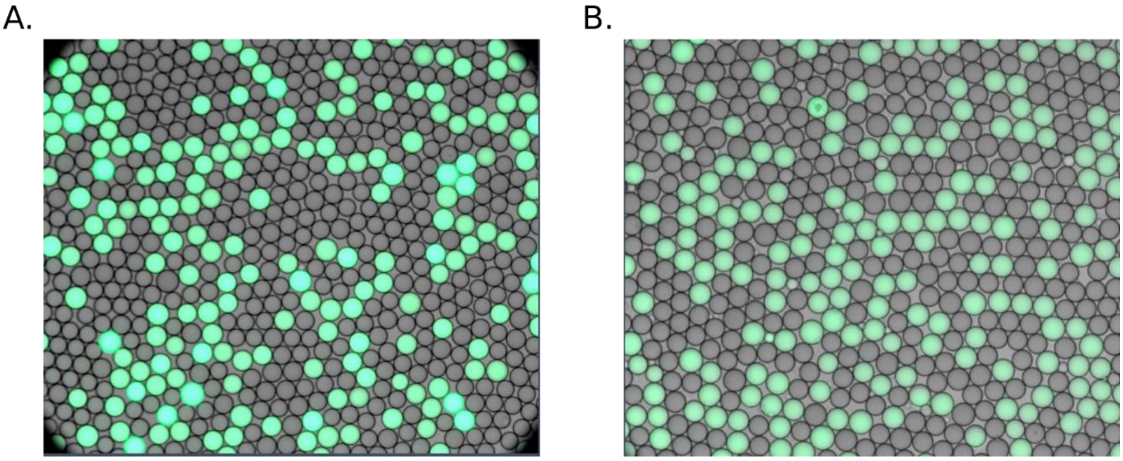

**Figure S1. Calcein leakage test in droplets.** Calcein crossing from one droplet to another or leaking to the surrounding oil would compromise the assay. To test for leakage, we generated droplets filled with calcein solution (70 mM in MOPS buffer) and analyzed them under a microscope immediately after generation (A). Then, droplets were transferred to a “droplet lung”[9] with homogeneous oxygen supply and incubated for 3 d at 30°C. After the incubation period, no leakage or cross move was observed within the droplet population (B).

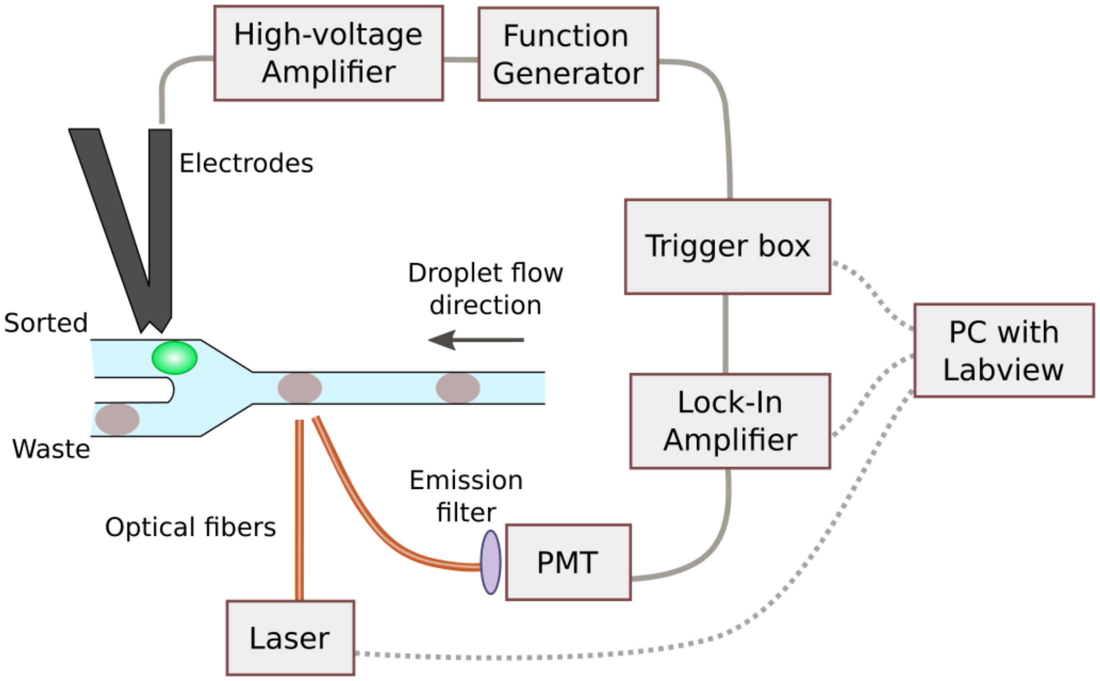

**Figure S2. Schematic of the droplet sorting setup**. Fluorescence signals from droplets are guided through optical fibers and detected by a PMT connected to a Lock-in Amplifier. The FPGA-based trigger box converts the analog voltage signal to a digital signal depending on sorting gates to generate a trigger signal. A function generator in conjugation with a high-voltage amplifier generates an alternating electric field in the electrodes of the chip for sorting droplets depending on the trigger signal. Sorting gates in the trigger box, laser, and lock-in amplifier are controlled using Labview-based software on a PC.[10]

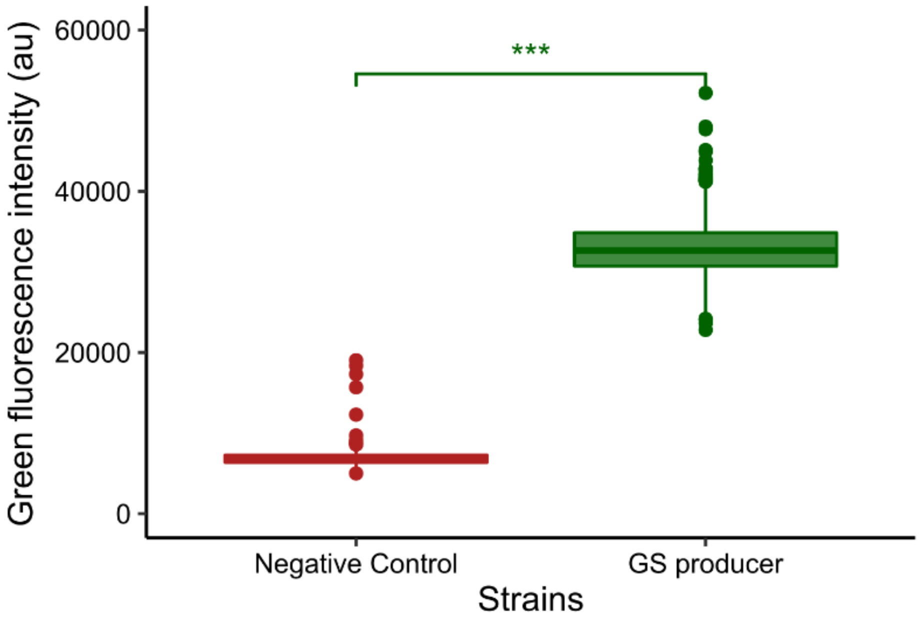

**Figure S3. Fluorescence signal in droplets.** To confirm that the signal resulting from the permeabilization of droplets is distinguishable from the signal measured from the negative control strain droplets, we co-incubated droplets with liposomes for 48 h at 28°C. The p-value obtained from performing a Student t-test indicates a highly significant difference between the two populations (p-value: 2.2E-16). Images in Figure 1D are obtained from the same experiment.

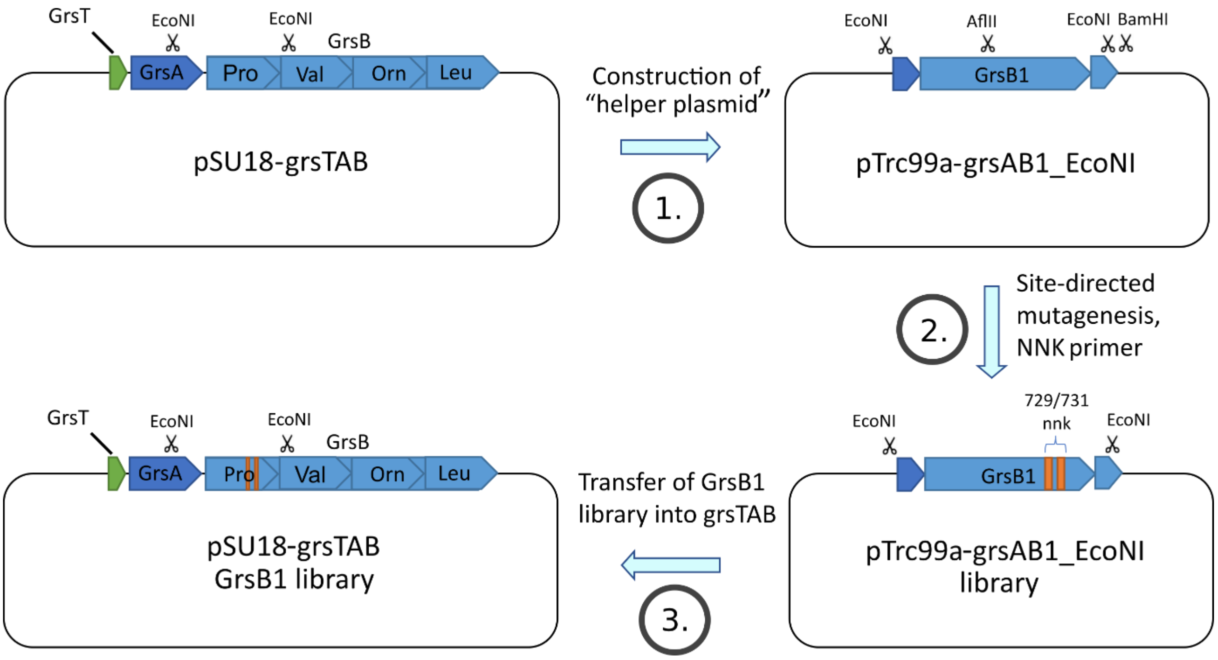

**Figure S4. Cloning of the GrsB1 library.** 1. Construction of pTrc99a-grsAB1_EcoNI, 2. Construction of pTrc99a-grsTAB_nnk helper plasmid, 3. Construction of pSU18-grsTAB_nnk Library.

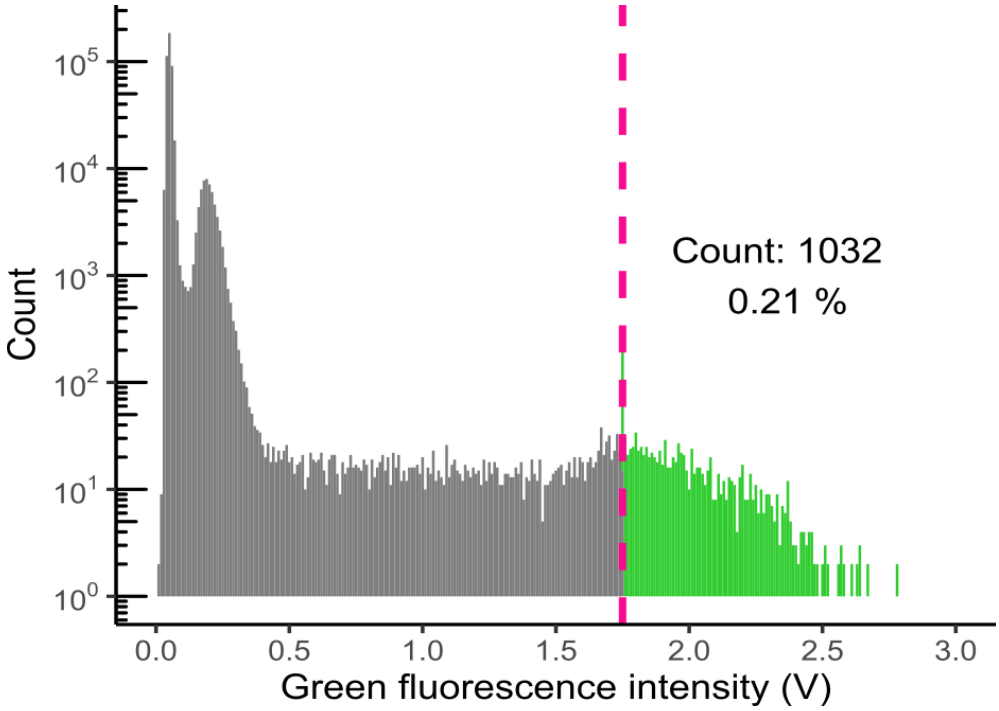

**Figure S5. Sorting statistics for the GrsB1 A-domain mutant library.** The sorting threshold (purple dashed line) was set at 1.75 relative fluorescence units. Fluorescent droplets surpassing the sorting threshold were collected. A parallel screening was performed with deactivated sorting gate.

### Supplementary Tables

**Table S1. List of primers.**

| Oligo | Sequence from 5' to 3' |
|---|---|
| pTrc99a-GrsB1_KpnI_fw_CTM | TGC GTG AAA CAT ATT TGC ACA ATG GGA GAA CAA TTA GTA GTT AAC AAT GAG |
| pTrc99a-GrsB1_KpnI_fw_CTN | TGC GTG AAA CAT ATT TGC ACA AAC GGA GAA CAA TTA GTA GTT AAC AAT GAG |
| pTrc99a-GrsB1_KpnI_fw_DTG | TGC GTG AAA CAT ATT GAT ACA GGC GGA GAA CAA TTA GTA GTT AAC AAT GAG |
| pTrc99a-GrsB1_KpnI_fw_ETS | TGC GTG AAA CAT ATT GAA ACA AGC GGA GAA CAA TTA GTA GTT AAC AAT GAG |
| pTrc99a-GrsB1_KpnI_fw_FTN | TGC GTG AAA CAT ATT TTT ACA AAC GGA GAA CAA TTA GTA GTT AAC AAT GAG |
| pTrc99a-GrsB1_KpnI_fw_GTF | TGC GTG AAA CAT ATT GGC ACA TTT GGA GAA CAA TTA GTA GTT AAC AAT GAG |
| pTrc99a-GrsB1_KpnI_fw_GTG | TGC GTG AAA CAT ATT GGC ACA GGC GGA GAA CAA TTA GTA GTT AAC AAT GAG |
| pTrc99a-GrsB1_KpnI_fw_GTH | TGC GTG AAA CAT ATT GGC ACA CAT GGA GAA CAA TTA GTA GTT AAC AAT GAG |
| pTrc99a-GrsB1_KpnI_fw_GTM | TGC GTG AAA CAT ATT GGC ACA ATG GGA GAA CAA TTA GTA GTT AAC AAT GAG |
| pTrc99a-GrsB1_KpnI_fw_HTM | TGC GTG AAA CAT ATT CAT ACA ATG GGA GAA CAA TTA GTA GTT AAC AAT GAG |
| pTrc99a-GrsB1_KpnI_fw_HTT | TGC GTG AAA CAT ATT CAT ACA ACC GGA GAA CAA TTA GTA GTT AAC AAT GAG |
| pTrc99a-GrsB1_KpnI_fw_KTF | TGC GTG AAA CAT ATT AAA ACA TTT GGA GAA CAA TTA GTA GTT AAC AAT GAG |
| pTrc99a-GrsB1_KpnI_fw_LTF | TGC GTG AAA CAT ATT CTG ACA TTT GGA GAA CAA TTA GTA GTT AAC AAT GAG |
| pTrc99a-GrsB1_KpnI_fw_LTG | TGC GTG AAA CAT ATT CTG ACA GGC GGA GAA CAA TTA GTA GTT AAC AAT GAG |
| pTrc99a-GrsB1_KpnI_fw_LTI | TGC GTG AAA CAT ATT CTG ACA ATT GGA GAA CAA TTA GTA GTT AAC AAT GAG |
| pTrc99a-GrsB1_KpnI_fw_LTW | TGC GTG AAA CAT ATT CTG ACA GAA GGA GAA CAA TTA GTA GTT AAC AAT GAG |
| pTrc99a-GrsB1_KpnI_fw_LTY | TGC GTG AAA CAT ATT CTG ACA TAT GGA GAA CAA TTA GTA GTT AAC AAT GAG |
| pTrc99a-GrsB1_KpnI_fw_MTG | TGC GTG AAA CAT ATT ATG ACA GGC GGA GAA CAA TTA GTA GTT AAC AAT GAG |
| pTrc99a-GrsB1_KpnI_fw_NTF | TGC GTG AAA CAT ATT AAC ACA TTT GGA GAA CAA TTA GTA GTT AAC AAT GAG |
| pTrc99a-GrsB1_KpnI_fw_NTV | TGC GTG AAA CAT ATT TGC ACA ATG GGA GAA CAA TTA GTA GTT AAC AAT GAG |
| pTrc99a-GrsB1_KpnI_fw_QTF | TGC GTG AAA CAT ATT CAG ACA TTT GGA GAA CAA TTA GTA GTT AAC AAT GAG |
| pTrc99a-GrsB1_KpnI_fw_RTA | TGC GTG AAA CAT ATT CGC ACA GCG GGA GAA CAA TTA GTA GTT AAC AAT GAG |
| pTrc99a-GrsB1_KpnI_fw_RTR | TGC GTG AAA CAT ATT CGC ACA CGC GGA GAA CAA TTA GTA GTT AAC AAT GAG |

| | |
|---|---|
| pTrc99a-GrsB1_KpnI_fw_RTV | TGC GTG AAA CAT ATT CGC ACA GTG GGA GAA CAA TTA GTA GTT AAC AAT GAG |
| pTrc99a-GrsB1_KpnI_fw_STM | TGC GTG AAA CAT ATT AGC ACA ATG GGA GAA CAA TTA GTA GTT AAC AAT GAG |
| pTrc99a-GrsB1_KpnI_fw_STN | TGC GTG AAA CAT ATT AGC ACA AAC GGA GAA CAA TTA GTA GTT AAC AAT GAG |
| pTrc99a-GrsB1_KpnI_fw_VTC | TGC GTG AAA CAT ATT GTG ACA TGC GGA GAA CAA TTA GTA GTT AAC AAT GAG |
| pTrc99a-GrsB1_KpnI_fw_VTG | TGC GTG AAA CAT ATT GTG ACA GGC GGA GAA CAA TTA GTA GTT AAC AAT GAG |
| pTrc99a-GrsB1_KpnI_fw_VTN | TGC GTG AAA CAT ATT GTG ACA AAC GGA GAA CAA TTA GTA GTT AAC AAT GAG |
| pTrc99a-GrsB1_KpnI_fw_VTV | TGC GTG AAA CAT ATT GTG ACA GTG GGA GAA CAA TTA GTA GTT AAC AAT GAG |
| pTrc99a-GrsB1_KpnI_fw_VTW | TGC GTG AAA CAT ATT GTG ACA GAA GGA GAA CAA TTA GTA GTT AAC AAT GAG |
| pTrc99a-GrsB1_KpnI_fw_WTK | TGC GTG AAA CAT ATT GAA ACA AAG GGA GAA CAA TTA GTA GTT AAC AAT GAG |
| pTrc99a-GrsB1_KpnI_fw_WTS | TGC GTG AAA CAT ATT TGG ACA AGC GGA GAA CAA TTA GTA GTT AAC AAT GAG |
| pTrc99a_GrsAB1_EcoNI_fw | ATT TCA CAC AGG AAA CTC GAG GCA GAA TAC CTA ACA AAG GAA TCG G |
| pTrc99a_GrsAB1_EcoNI_rev | TGG TGA TGA GAT CTG GAT CCT TGA ATG CCT AAC GTA GGA AGT TCC |
| GrsAB1_grsTAB_fw | TAC GCA GAA TAC CTA ACA AAG GAA TCG G |
| GrsAB1_grsTAB_rev | TTA TAT TGA ATG CCT AAC GTA GGA AGT TCC |
| GrsB1-N-seq-4 | GTA AAA CGT GAA AAT ATT GAA GTA TTA TCC |
| GrsB1_H765_rev | ATT GTG TAA ATG TAC GTT ATG TTC ATG C |
| pSU18-GrsB1_V969KpnI_fw | CAA ACG CAA AAT ATG TGG TAC CTA CAA ATG AGC TGG AAG AAA AAT TGG |
| pSU18-GrsB1_V969_rev | ACA TAT TTT GCG TTT GTA TTC ACA ATC C |
| GrsB1_AflII_fw | TTA GCC AGA TTC TTA AGA GAA AAA GGC |
| GrsB1_I729nnk_A731nnk_fw | TGC GTG AAA CAT ATT nnk ACA nnk GGA GAA CAA TTA GTA GTT AAC AAT GAG |

**Table S2. Sequence of protein GrsB1.***

```
MSTFKKEHVQDMYRLSPMQEGMLFHALLDKDKNAHLVQMSIAIEGIVDVELLSESLNILIDRYDVFRT
TFLHEKIKQPLQVVLKERPVQLQFKDISSLDEEKREQAIEQYKYQDGETVFDLTRDPLMRVAIFQTGK
VNYQMIWSFHHILMDGWCFNIIFNDLFNIYLSLKEKKPLQLEAVQPYKQFIKWLEKQDKQEALRYWKE
HLMNYDQSVTLPKKKAAINNTTYEPAQFRFAFDKVLTQQLLRIANQSQVTLNIVFQTIWGIVLQKYNS
TNDVVYGSVVSGRPSEISGIEKMVGLFINTLPLRIQTQKDQSFIELVKTVHQNVLFSQQHEYFPLYEI
QNHTELKQNLIDHIMVIENYPLVEELQKNSIMQKVGFTVRDVKMFEPTNYDMTVMVLPRDEISVRLDY
NAAVYDIDFIKKIEGHMKEVALCVANNPHVLVQDVPLLTKQEKQHLLVELHDSITEYPDKTIHQLFTE
QVEKTPEHVAVVFEDEKVTYRELHERSNQLARFLREKGVKKESIIGIMMERSVEMIVGILGILKAGGA
FVPIDPEYPKERIGYMLDSVRLVLTQRHLKDKFAFTKETIVIEDPSISHELTEEIDYINESEDLFYII
YTSGTTGKPKGVMLEHKNIVNLLHFTFEKTNINFSDKVLQYTTCSFDVCYQEIFSTLLSGGQLYLIRK
ETQRDVEQLFDLVKRENIEVLSFPVAFLKFIFNEREFINRFPTCVKHIITAGEQLVVNNEFKRYLHEH
NVHLHNHYGPSETHVVTTYTINPEAEIPELPPIGKPISNTWIYILDQEQQLQPQGIVGELYISGANVG
RGYLNNQELTAEKFFADPFRPNERMYRTGDLARWLPDGNIEFLGRADHQVKIRGHRIELGEIEAQLLN
CKGVKEAVVIDKADDKGGKYLCAYVVMEVEVNDSELREYLGKALPDYMIPSFFVPLDQLPLTPNGKID
RKSLPNLEGIVNTNAKYVVPTNELEEKLAKIWEEVLGISQIGIQDNFFSLGGHSLKAITLISRMNKEC
NVDIPLRLLFEAPTIQEISNYINGGSRS*
```

*The mutated positions are highlighted in red.

**Table S3. Specificity codes of Pro-A-domains.**[#]

| | | Sequence position* | | | | | | | | |
|---|---|---|---|---|---|---|---|---|---|---|
| | | 235 | 236 | 239 | 278 | 299 | 301 | 322 | 330 | 331 |
| **NRPS** | **UniProt ID** | 659 | 660 | 663 | 702 | 729 | 731 | 755 | 763 | 764 |
| Gramicidin S | P0C064 | D | V | Q | S | I | A | H | V | V |
| Virginiamycin S | O05647 | D | V | Q | Y | A | A | H | V | M |
| Pristinamycin I | O07944 | D | V | Q | Y | A | A | H | V | M |
| Tyrocidine | O30408 | D | V | Q | S | I | A | H | V | V |
| Fengycin | O30980 | D | V | Q | V | I | A | H | V | V |
| D-lysergyl-peptide | O94205 | D | I | T | L | V | A | G | L | I |
| Plipastatin | P94459 | D | V | Q | F | I | A | H | V | V |
| HC toxin | Q01886 | D | I | A | V | I | T | V | L | I |
| Iturin A | Q93I55 | D | V | Q | F | I | A | H | V | V |
| Syringopeptin | Q9FDB3 | D | V | Q | Y | I | A | H | V | V |
| Mycosubtilin | Q9R9J0 | D | V | Q | F | I | A | H | V | V |
| Peptaibol | Q8NJX1 | D | V | L | F | C | G | L | I | C |

[#]Sequences have been retrieved from the supplementary of ref. [14]. The positions mutated in this work are highlighted in red.

*Top row: sequence position in GrsA (PDB code 1amu). Bottom row: sequence position in GrsB (first entry).

**Table S4. Enrichment and catalytic activity of selected GrsB1 variants.**†

| Protein sequences | Enrichment factor# | Protein yield$ [mg] | $k_{cat}/K_M$§ [$mM^{-1}$ $min^{-1}$] | ProHA* [µM] |
|---|---|---|---|---|
| ITA (WT) | 1.00 | 36.0 | 135 | 17.6 |
| **CTM** | 11.3 | 30.7 | 1.19 | 0.14 |
| **CTN** | 8.84 | 35.4 | 0.234 | 6.93 |
| **GTH** | 7.34 | 39.9 | 1.97 | 0.39 |
| **NTF** | 6.49 | 41.9 | 2.19 | 0.57 |
| **ETS** | 3.73 | 34.4 | 0.461 | 10.0 |
| **WTS** | 2.08 | 24.8 | 32.3 | 0.79 |
| **KTF** | 1.69 | 42.2 | 0.294 | 6.29 |
| **LTY** | 1.47 | 27.0 | 0.760 | 0.73 |
| **GTM** | 1.21 | 9.2 | 0.128 | 0.07 |
| **GTF** | 1.01 | 24.9 | 0.553 | 6.90 |
| **HTM** | 0.83 | 19.7 | 0.409 | 2.16 |
| **VTN** | 0.80 | 30.9 | 0.058 | 2.02 |
| **QTF** | 0.55 | 7.6 | 1.32 | 1.09 |
| **STN** | 0.51 | 25.7 | 0.160 | 7.33 |
| **RTA** | 0.42 | 26.5 | 1.93 | 4.91 |
| **VTW** | 0.34 | 9.5 | 0.283 | 0.81 |
| **STM** | 0.25 | 25.2 | 1.05 | 0.27 |
| **LTW** | 0.21 | 20.9 | 0.276 | 0.80 |
| **HTT** | 0.18 | 26.5 | 0.120 | 0.34 |
| **VTC** | 0.078 | 16.6 | 0.113 | 2.72 |
| **LTF** | 0.060 | 31.3 | 2.50 | 7.08 |
| **FTN** | 0.026 | 12.0 | 0.027 | 1.26 |
| **VTV** | 0.014 | 16.8 | 0.167 | 0.20 |
| **NTV** | 0.0090 | 2.8 | 0.406 | 0.01 |
| **MTG** | 0.0090 | 32.4 | 0.296 | 0.35 |
| **LTG** | 0.0077 | 16.3 | 0.163 | 0.58 |
| **GTG** | 0.0070 | 31.0 | 0.064 | 0.18 |
| **VTG** | 0.0067 | 7.0 | 0.069 | 3.81 |
| **DTG** | 0.0021 | 3.2 | 0.023 | 0.01 |
| **RTV** | 0.00024 | 25.5 | ND | ND |
| **RTR** | 0.00012 | 0.0 | NA | NA |
| **WTK** | 0.00008 | 0.0 | NA | NA |
| **LTI** | 0.00003 | 36.2 | 0.113 | ND |

†NA: not applicable as no protein was obtained. ND: not detectable.

#Enrichment factor calculated by dividing the frequency of a mutation observed in each population (sorted population vs. the unsorted population) compared to the wild-type.

$$Enrichment\ factor = \frac{\frac{variant\ frequency\ in\ sorted\ population}{variant\ frequency\ in\ unsorted\ population}}{\frac{wildtype\ frequency\ in\ sorted\ population}{wildtype\ frequency\ in\ unsorted\ population}}$$

$Yield of heterologously expressed and purified protein.

$^{§}k_{cat}/K_M$ [$mM^{-1}$ $min^{-1}$] was measured by saturation kinetics of hydroxylamine stimulated pyrophosphate release with Pro as a substrate.

*ProHA was measured using the HAMA assay.[6]
